# Somatic mutation in human cerebellum illustrates neuron type-specific patterns of age-related mutation

**DOI:** 10.64898/2026.02.27.708647

**Authors:** Kow Essuman, Yingxi Yang, Eitan Goodman, Christie N. Cambridge, Chunhui Cai, Zheming An, Shulin Mao, Monica Devi Manam, Benjamin Finander, Sattar Khoshkhoo, Liang Sun, August Yue Huang, Christopher A. Walsh

## Abstract

Human neurodegenerative disorders are characterized by exquisite specificity for neuronal types, but the basis of this is unknown. Here, we show that cerebellar granule neurons (GN)—the most abundant neuronal type in the human brain—accumulate somatic mutations in patterns highly distinct from cerebral cortical neurons, and more closely resembling oligodendroglia and other dividing cells. We find shared mutational signatures between normal aging GNs and medulloblastoma subtypes, suggesting the GN lineage as a tumor cell of origin. Whole-genome sequence of multiple single GNs from the same donor allowed analysis of specific times of neurogenesis, revealing a rich lineage tree that includes GNs that become postmitotic 2 years or more after birth, yet migrating postnatally to populate both the cerebellar vermis and the distant cerebellar hemisphere. Our results show that neuronal type-specific somatic mutation patterns enlighten normal development, cancer origins and potentially the cell type-specificity of neurodegeneration.

## Main Text

The human brain is estimated to contain as many as one thousand distinct neuronal types, defined by their morphology, connections, physiological activity, transcriptomic diversity (*1*, *2*), and prominently the fact that they are postmitotic and do not undergo further cell division once formed (*3*). Despite their postmitotic state, cerebral cortical neurons show somatic mutations that reflect their cell lineage during development (*4*, *5*), but also continue to accumulate mutations throughout their postmitotic life (*6–8*), at rates comparable to those of dividing cells such as hematopoietic stem cells (*6*, *9*, *10*). In cerebral cortical pyramidal neurons, where they have been studied, these age-related mutations appear to be related to DNA damage occurring in relation to gene transcription, since they are enriched in genes and regulatory sequences (*5*, *7*, *8*, *11*), in contrast to somatic mutations in oligodendrocytes (OLs), which show little relation to transcription and a stronger relationship to cell cycle-related DNA replication (*11*).

Despite the potential importance of age-related somatic mutations in neurons for neurodegenerative disease (*12–14*), they have so far only been intensively studied in a single neuronal type, the excitatory pyramidal neurons of the cerebral cortex (*12*, *13*). Therefore, it is unknown how the rules defined for cortical neurons may or may not apply to other neuronal types in the central nervous system. Degenerative diseases affecting the nervous system are as rule quite specific for cell types (*15*) emphasizing the need to study somatic mutation in other neuronal types.

Granule neurons (GNs) of the cerebellum are the most numerous neurons of the human brain, estimated to be more abundant than all other neurons in the brain combined, and have several distinctive features (*16*, *17*). They have small nuclei and perikarya, and are formed relatively late in development, with animal studies suggesting that they continue to be formed for some time after birth (*16–18*). They derive from precursors of the rhombic lip, which migrate out tangentially over the outer surface of the incipient cerebellum, forming a cell-dense layer referred to as the external granule cell layer (EGL) (*16*, *17*, *19*). Classical histological studies in humans show that this EGL persists at least a year after birth (*20–23*), though the extent of formation of neurons after birth in human cerebellum is not known.

Most degenerative disorders that affect the pyramidal neurons of the cerebral cortex do not typically affect GNs (*15*). However, there are GN-specific forms of degeneration, and disorders of cerebellar development and cancer (*24–26*) that prompt a need for better understanding of the somatic mutation landscape in the cerebellum. Our analysis of age-related somatic mutation rates and patterns in GNs show widespread differences from cortical neurons and illustrate the unique development of the human cerebellum.

### Cerebellar GNs accumulate somatic mutations at distinct rates from cortical neurons

We first purified NeuN+ nuclei from post-mortem human cerebellar hemisphere or vermis using fluorescent activated nuclei sorting (FANS) (Fig. 1A and fig. S1A). NeuN+ was chosen to enrich for neuronal populations, but to ensure our sorting strategy yielded mature GNs rather than their precursors, we performed single-nuclei RNA sequencing (snRNA-seq) on the sorted population (Fig. 1, A to C, and fig. S1B). We generated 38,223 nuclei from eight samples across four neurotypical individuals aged 19.8, 42.2, 59, 82.7 years (Fig. 1B, fig. S1B, table S1). We found that high quality cells from snRNA-seq yielded 99% GN purity with marker genes including *GRIK2* and *GABRA6* (*27*, *28*) to confirm mature GN purity (Fig. 1, B and C). Less than 1% of sorted neurons lacked expression of *GABRA6* and these cells were classified as differentiating, immature GNs (Fig. 1, B and C, and fig. S1B).

**Fig. 1.**
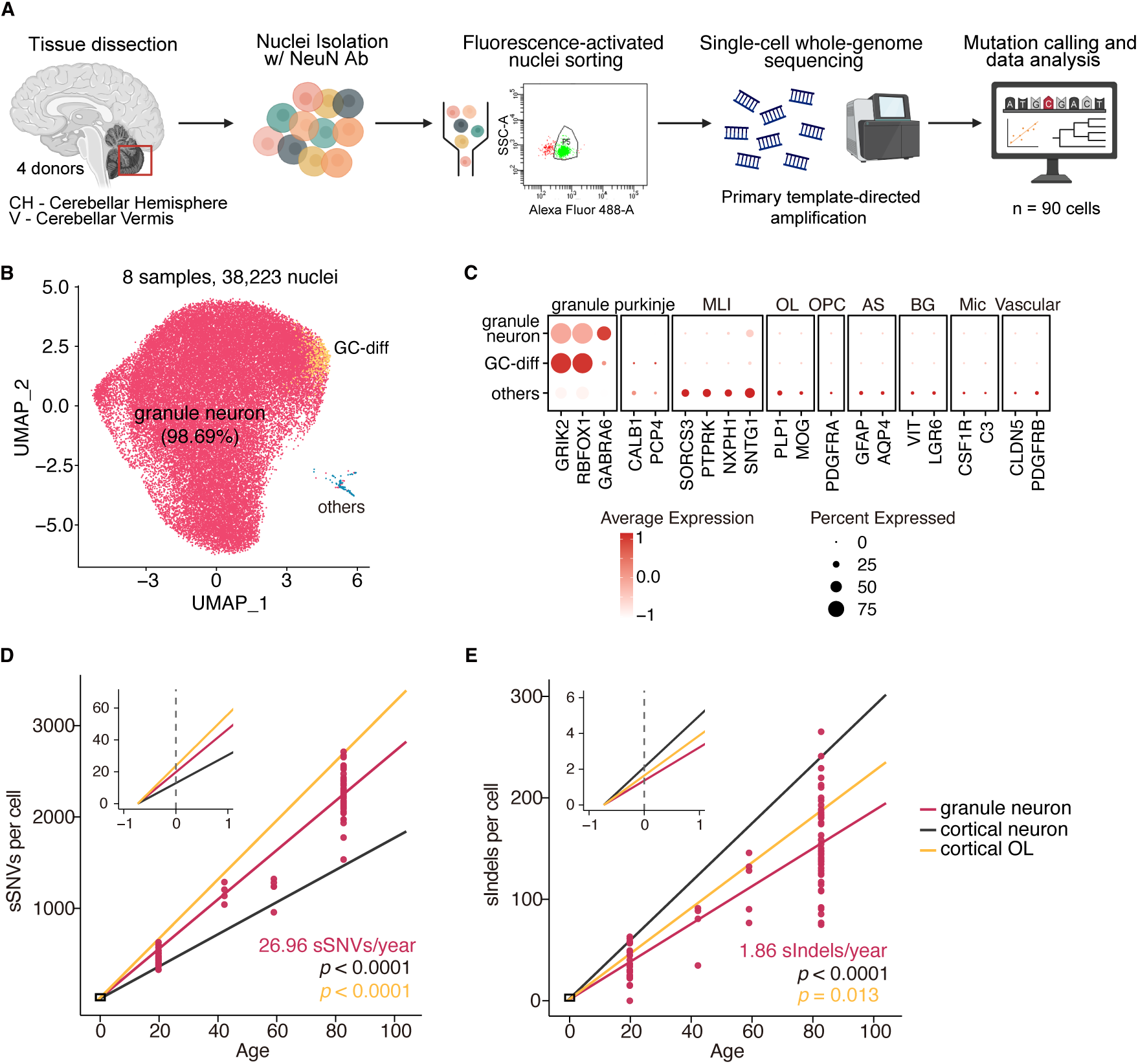
Genome-wide rates of somatic mutation accumulation in cerebellar GNs. A. Schematic of experimental design. GNs were isolated from postmortem human cerebellar tissue, followed by primary template amplification (PTA), whole-genome sequencing, and downstream variant analysis. B. Uniform manifold approximation and projection (UMAP) representation of snRNA-seq profiles generated from NeuN-sorted cerebellar samples. Major cell types and their relative proportions are labeled. GC-diff, granule cell-differentiating cells. C. Dotplot showing the mean expression of marker genes and the percentage of cells expressing them for each annotated cell type. MLI, molecular layer interneurons; OL, oligodendrocyte; OPC, oligodendrocyte precursor cell; AS, astrocyte; BG, Bergmann glia; Mic, microglia. D. sSNV burden as a function of age in GNs (red), cortical neurons (black), and cortical OLs (orange). Data points represent single GNs; trend lines reflect linear mixed-effects model; the inset shows an enlarged view of modeled early-life accumulation (fertilization to one year). P values compare GNs with cortical neurons and OLs (GN vs cortical neuron: *p* < 0.0001; GN vs cortical OL: *p* < 0.0001). E. sIndel burden analyzed as in (D). GN vs cortical neuron: *p* < 0.0001; GN vs cortical OL: *p* = 0.013.

To determine how somatic mutations accumulate with age in GNs, we performed single-cell whole genome amplification with primary template amplification (PTA) as previously described (*8*, *12*), and performed whole genome sequencing (scWGS) on 90 single GNs from the four individuals (age 19.8, 42.2, 59, 82.7), targeting 30X coverage for each cell (range 19–63X coverage) (Fig. 1A). We then analyzed single genomes using our well established SCAN2 genotyper (*8*, *11*) to accurately identify somatic single nucleotide variants (sSNVs) and somatic insertions-deletions (sIndels) from each cell (table S2 and S3). Somatic mutations in GNs were then compared with those in 56 cerebral cortical pyramidal neurons and 66 OLs previously sequenced from human prefrontal cortex (*8*, *11*) (table S1).

Similar to cerebral cortical neurons and OLs, cerebellar GNs accumulate somatic sSNVs and sIndels with aging. The rate of sSNV accumulation in GNs is 26.96 sSNV/yr (95% confidence interval [CI]: 25.1–29.0), which is faster than that of cortical neurons 17.59 sSNVs/yr (CI: 16.1–19.1; for the difference, p = 3.9 × 10^−10^, Tukey-corrected t-test), but slower than cortical OLs 32.32 sSNVs/yr (CI: 30.6–34.0; p = 1.3 × 10^−7^, Tukey-corrected t-test) (Fig. 1D). In contrast, sIndels accumulate at a slower rate in GNs (1.86 sIndels/yr, CI: 1.6–2.1) compared to both cortical neurons (2.88 sIndels/yr, CI: 2.7–3.1; p = 4.5 × 10^−9^, Tukey-corrected t-test), and cortical OLs (2.24 sIndels/yr, CI: 2.0–2.5; p = 0.013, Tukey-corrected t-test) (Fig. 1E). These trends persisted after controlling for multiple quality metrics (fig. S2). Analysis of mutation burden and mutational spectra across cerebellar regions revealed no marked differences between GNs isolated from the midline vermis region, or the more laterally situated, functionally distinct, cerebellar hemispheres (fig. S3). Altogether, these findings suggest that different cell types in the human brain, even among neurons, accumulate somatic mutations at different rates, and hence may have unique vulnerabilities or response mechanisms to genomic damage.

### sSNV signatures of cerebellar GNs differ from that of cortical neurons

To better understand the potential mutational processes occurring with age and to provide clues as to their potential underlying mechanisms, we performed mutational spectrum analysis of both sSNVs and sIndels of GNs. GNs showed a significant increase in C>A (p = 5.3 × 10^−21^, two-tailed Wilcoxon test) and C>T (p = 1.9 × 10^−10^, two-tailed Wilcoxon test) mutations compared to cortical neurons (Fig. 2A). Notably, comparison of mutational spectra indicated that the mutational spectrum of GNs was more similar to OLs (cosine similarity 0.973) than to cortical neurons (cosine similarity 0.896) (Fig. 2, A and B).

**Fig. 2.**
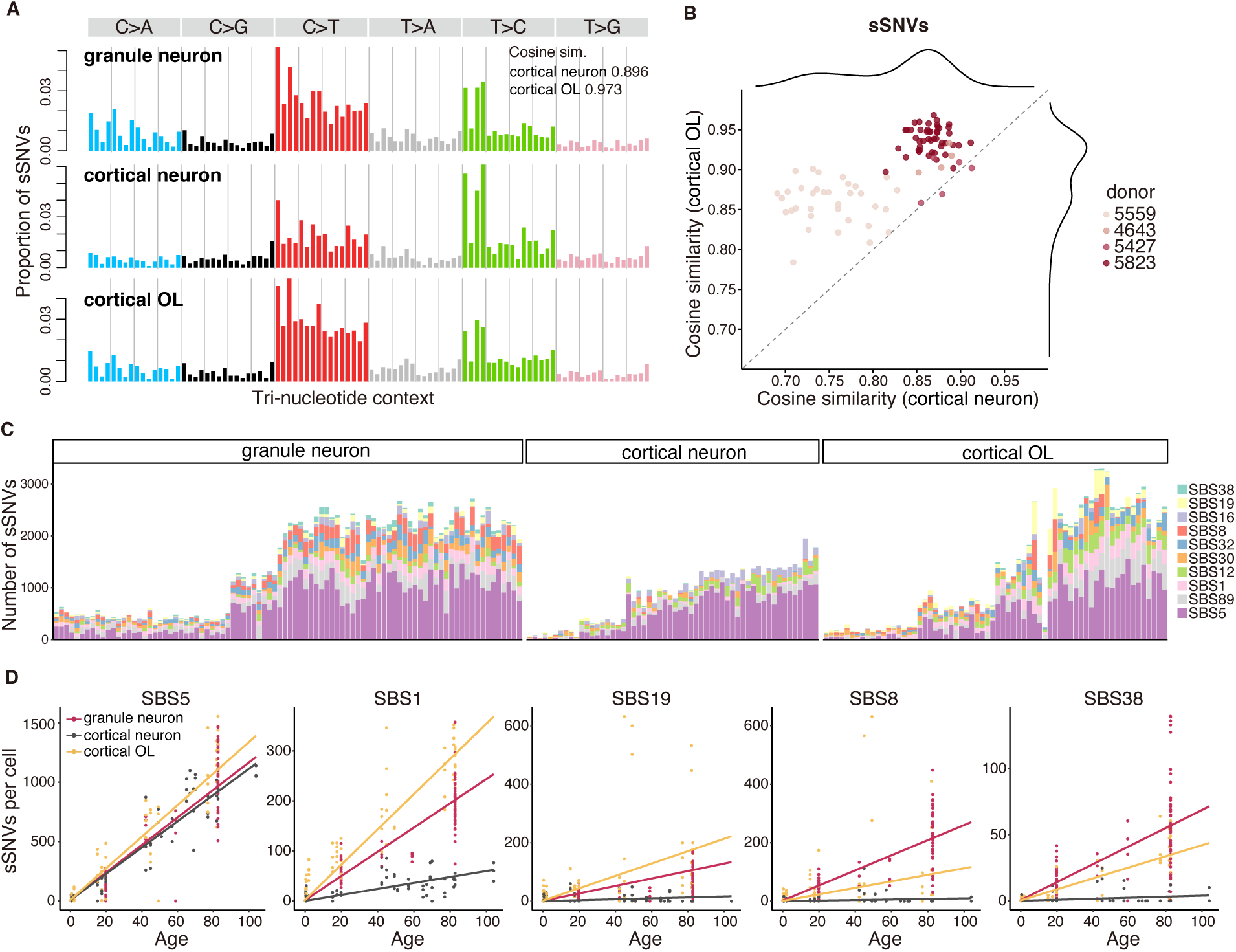
Cerebellar GNs show stronger age-associated increases in cell-division-related mutational signatures than cortical neurons. A. sSNV mutational spectrum of the three cell types, with cosine similarities shown for GNs relative to cortical neurons and OLs. B. Cosine similarity of sSNV spectrum from each GN relative to cortical neurons and OLs. Points represent single GNs colored by subject. C. Decomposition of sSNV burden into COSMIC SBS signatures in GNs, cortical neurons, and OLs. Subjects are ordered by increasing age from left to right. D. Contributions of the selected COSMIC SBS signatures to the sSNV burden in GNs, cortical neurons, and OLs.

To disentangle the underlying molecular processes responsible for the accumulation of mutations in GNs, we performed COSMIC signature decomposition analysis as previously described (*29*), combining mutational patterns refitting with stepwise regression to prioritize the best-fitting signatures (*30*). This revealed an unexpected accumulation of the cell-division-associated mutational signature SBS1 in aging GNs (Fig. 2, C and D). SBS1 is enriched in rapidly dividing cells including cancer cells and is thought to reflect enzymatic or spontaneous deamination of 5-methylcytosine to thymine during the cell cycle, and is enriched at CpG islands in the genome (*6*, *31*, *32*). We found GNs from the aged individual showed markedly elevated SBS1 exposure relative to younger individuals, despite the postmitotic nature of GNs. On the other hand, age-matched cortical neurons do not show such high rates of SBS1 accumulation (Fig. 2, C and D; for the difference between GNs and cortical neurons, p = 2.7 × 10^−14^, Tukey-corrected t-test). The rate of SBS1 accumulation in GNs however appears to be slower than that in OLs (Fig. 2, C and D; p = 9.7 × 10^−13^, Tukey-corrected t-test). Other mutational signature patterns commonly seen in dividing cells also increased with age in GNs, including SBS19 (for the difference between GNs and cortical neurons, p = 2.5 × 10^−3^; GNs and OLs, p = 0.01; Fig. 2D) and SBS32 (GNs and cortical neurons, p = 4.4 × 10^−7^; GNs and OLs, p = 0.27, Tukey-corrected t-test; fig. S4B), observed in blood, liver and brain cancers (*33–36*). Our data therefore reveals shared mutational signatures and patterns between GNs and OLs, which suggests shared underlying mechanisms despite their differences in cell type and lineage.

The aging-associated SBS5 signature accumulated in cerebellar GNs at rates comparable to the rate of SBS5 accumulation in cortical neurons and OLs (Fig. 2D). SBS5 is a “clock-like” mutational signature that accumulates in all cancers, cells, and tissues studied to date, and is the major driver of age-related mutation accumulation in cerebral cortical neurons (*11*), suggesting that it accumulates independently of cell division. SBS16, a transcription-associated mutational signature that also increases in cortical neurons with age (*11*), accumulated at a slower rate in GNs (p = 3.5 × 10^−4^, Tukey-corrected t-test; fig. S4). The higher burden of C to A mutations in GNs compared to cortical neurons or OLs is likely due to SBS8 and SBS38 signature mechanisms (Fig. 2, C and D). The etiology or underlying mechanisms of these two signatures remain to be fully elucidated, though SBS8 has been linked to deficient nucleotide excision repair of oxidative lesions (*37*, *38*). SBS38 has been linked to ultraviolet light-associated melanoma (*34*), but the mechanism of its accumulation in GNs is unclear.

### sIndel signatures of cerebellar GNs differs from cerebral cortical neurons

Analogous to our sSNV analysis, GNs accumulate sIndels with signatures more closely resembling OLs (cosine similarity 0.978) than cortical neurons (cosine similarity 0.773) (Fig. 3, A and B). GNs show a lower number of 1 base-pair insertions of thymine nucleotides compared to cortical neurons (p = 2.0 × 10^−9^, two-tailed Wilcoxon test; Fig. 3A). Further, ID4 accumulates at a much slower rate in aging GNs than in cortical neurons (p = 4.9 × 10^−10^, Tukey-corrected t-test; Fig. 3, C and D). ID4-like signatures have recently been identified as a hallmark of several neurodegenerative disorders (*13*, *39*), and appear to reflect Topoisomerase 1 (TOP1)-mediated DNA damage (*39*, *40*). ID2 and ID9 signatures on the other hand, accumulate at higher rates in GNs than cortical neurons (ID2: p = 4.8 × 10^−2^, ID9: p = 2.1 × 10^−2^), but at similar rates to OLs (ID2: p = 0.46, ID9: p = 0.99, Tukey-corrected t-test; Fig. 3, C and D). While the etiology of ID9 is unknown, ID2 is frequently observed in cancer tissues and is the result of DNA damage induced by replication slippage (*34*). In all, the sIndel data are consistent with our sSNV data supporting the idea that the mutational forces and mechanisms in cerebellar GNs are more similar to those in OLs than to cerebral cortical neurons.

**Fig. 3.**
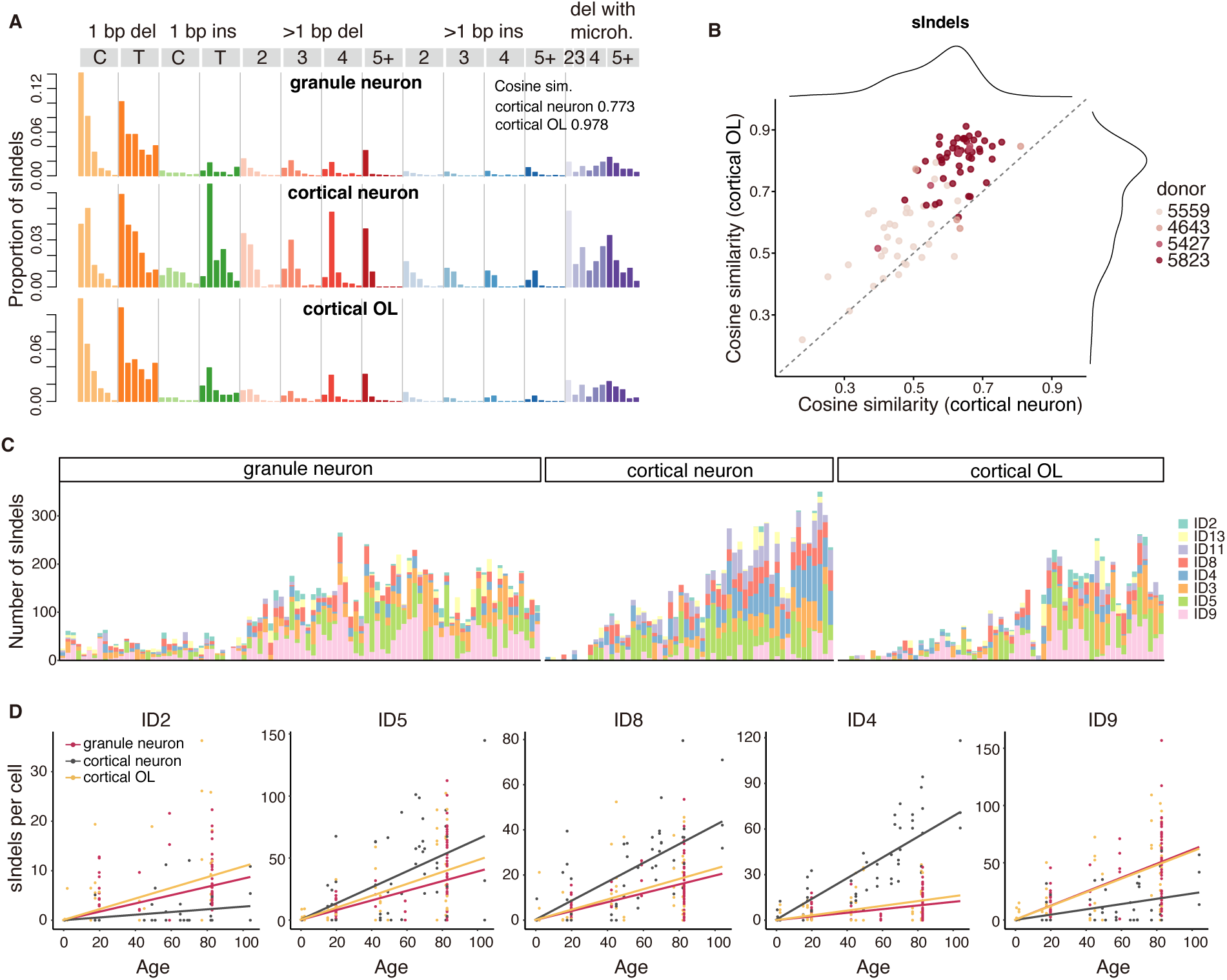
sIndel spectra of GNs show greater similarity to cortical OLs than to cortical neurons. A. sIndel mutational spectrum of the three cell types, with cosine similarities shown for GNs relative to cortical neurons and OLs. B. Cosine similarity of sIndel spectrum from each GN relative to cortical neurons and OLs. Points represent single GNs colored by subject. C. Decomposition of sIndel burden into COSMIC indel signatures in GNs, cortical neurons, and OLs. Subjects are ordered by increasing age from left to right. D. Contributions of the selected COSMIC indel signatures to the sIndel burden in GNs, cortical neurons, and OLs.

### Genomic distribution and functional impact of GN somatic mutation

Patterns of mutational enrichment in GNs also more closely resembled mutational patterns in OLs than cortical neurons (Fig. 4, A to D). Cortical neurons show enrichment of sSNVs and sIndels in genic regions, whereas oligodendrocyte mutations are enriched in intergenic regions (*11*). GNs show enrichment of sSNV and sIndels in intergenic regions similar to OLs but distinct from cortical neurons (Fig. 4, A to D). Specifically, sSNVs were enriched in intergenic regions but depleted in exonic and intronic genic regions (Fig. 4, A and B). Similar patterns of intergenic enrichment were observed with GN sIndels (Fig. 4, C and D). Consistent with this pattern, GNs and OLs had fewer moderate-impact sSNVs than cortical neurons (Fig. 4E). Intriguingly, unlike OLs, GNs harbored a significantly higher fraction of high-impact sIndels largely localized in exonic regions (Fig. 4F and fig. S5A). The genes predicted to be affected are involved in metabolism-related pathways, including carbohydrate metabolism as well as neurotransmitter activity (Fig. 4G). Future studies should clarify the functional effect of these predictions.

**Fig. 4.**
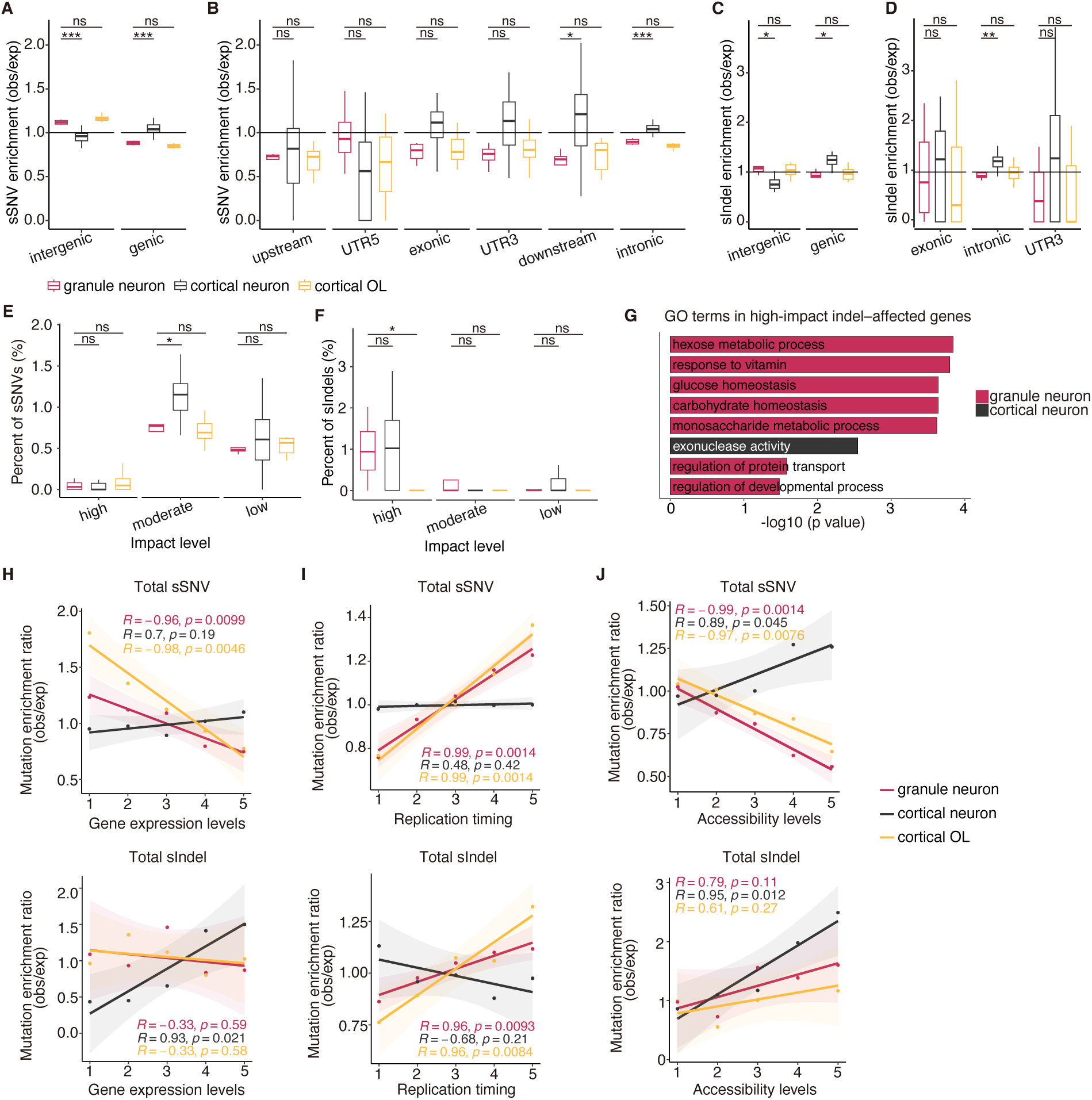
Enrichment analysis of sSNVs and sIndels in GNs. A–D. Genomic enrichment patterns of sSNVs (A, B) and sIndels (C, D) across cell types, displayed for intergenic and genic regions (A, C) and for detailed genic annotations (B, D). obs, observed; exp, expected (1,000 permutations). Significance denoted as: ns p ≥ 0.05, *p < 0.05, **p < 0.01, ***p < 0.001, ****p < 0.0001 using a two-tailed Wilcoxon test. E–F. Percentages of sSNVs (E) and sIndels (F) predicted as high-, moderate-, or low-impact in GNs, cortical neurons, and OLs. P value of two-tailed Wilcoxon test is provided. G. Gene ontology (GO) terms significantly enriched in genes affected by high-impact indels in granule and cortical neurons. H–J. sSNV and sIndel enrichment in relation to gene expression (H), replication timing (I), and chromatin accessibility (J). Gene expression and chromatin accessibility profiles for GNs were obtained from our current study and Ament et al.(*28*), respectively, whereas profiles for cortical neurons and OLs were taken from Jeffries et al. (*49*) and Ganz et al. (*11*). Replication timing was derived from ENCODE Repli-seq of 15 cell lines. Expected mutation density was computed from 1,000 permutations; points show the mean observed/expected ratio across subjects, and lines represent linear fits with 95% confidence intervals. Pearson’s correlation coefficient and P value are provided.

The density of sSNVs is positively associated with transcription levels in cortical neurons but negatively associated with transcriptional levels in OLs (*8*, *11*). Integrating our snRNA-seq data revealed negative correlations between GN sSNVs and gene expression, mirroring the pattern in OLs (Fig. 4H). sIndels followed a similar negative trend although this was not statistically significant (Fig. 4H). Leveraging replication timing profiles from 15 ENCODE Repli-seq cell lines, we found that similar to OLs, GNs showed an enrichment of sSNVs and sIndels in late-replicating genomic regions, an area of the genome thought to be transcriptionally silent (Fig. 4I). In line with their depletion in highly expressed genes, our GN sSNVs were also depleted in more accessible chromatin regions in a published dataset on human cerebellum (*28*) (Fig. 4J). This pattern matches that of OLs but again differs from cortical neurons (Fig. 4J). sIndels did not show a significant correlation with chromatin accessibility (Fig. 4J). Additionally, by dissecting mutational signature-specific enrichment patterns, we found SBS1, SBS8 and SBS38 contributing to sSNV enrichment in low-expression genes, late-replicating regions, and inaccessible genomic regions (fig. S5). Taken together, our data suggest that somatic mutations in GNs are enriched in inactive and less-highly transcribed regions of the genome, and that certain sIndels could have damaging effects affecting cellular metabolism.

### Lineage analysis of sorted single cerebellar GNs

Cerebellar GNs continue to be produced from precursor cells for some time after birth in animal models (*17*, *18*), and the persistence of the EGL in the immediate postnatal period in humans (*16*) suggests postnatal GN generation as well, prompting us to examine cell lineage and the timing of GN production directly in human brain. We focused on our oldest subject (age 82.7 years) where we had generated 47 GNs sequenced at an average of 30X. To expand the number of cells and strengthen our analysis, we isolated and whole-genome sequenced an additional 131 GNs at an average of 10X coverage, a depth sufficient for accurate sSNV calling, supported by identical mutational spectra between 30X and 10X cells from the same individual (cosine similarity 0.992) (fig. S6, A and B). Further, as our SCAN2-based mutation calling may fail to detect some early clonal mutations present in the bulk control samples, we additionally generated 250X bulk WGS on matched cerebellum samples and called clonal mutations using MosaicForecast (*41*) to allow capture of additional clonal mutations (fig. S6B). Shared sSNVs identified clonally related neurons, while the count of unshared somatic sSNVs, since they accumulate in a highly linear, clock-like fashion with respect to postnatal age, could be used to approximate the age of the donor at the time of the most recent common ancestor (MRCA) for each neuronal lineage (*11*) by dividing genome-wide private sSNVs by the annual accumulation rate from GN aging trend line, and subtracting from the donor age at death following model-based offset correction (see Methods).

These clonal analyses reveal three major clades (a, b, c), each defined by distinct sets of clonal sSNVs, giving rise to the 178 GNs after fertilization (Fig. 5A and fig. S6C). We found that most ancestral clones across all three clades arose postnatally, with 45 out of 47 MRCA events dated after birth (Fig. 5, A and B), consistent with extensive GN formation postnatally. Notably, within clades b and c, the latest MRCAs for sub-clades b’ and c’ emerged as late as approximately two years of age (1.95 years, 95% CI: 1.76–2.15; Fig. 5, A and B). Moreover, we find that there is extensive spatial intermingling of clones that ultimately define the cerebellar hemisphere (CH) and vermis (V) (Fig. 5C), with cells from both CH and V being dispersed through all three clades a, b, c, and persisting even within those late-born sub-clades (a’, b’, c’) (Fig. 5, A to E).

**Fig. 5.**
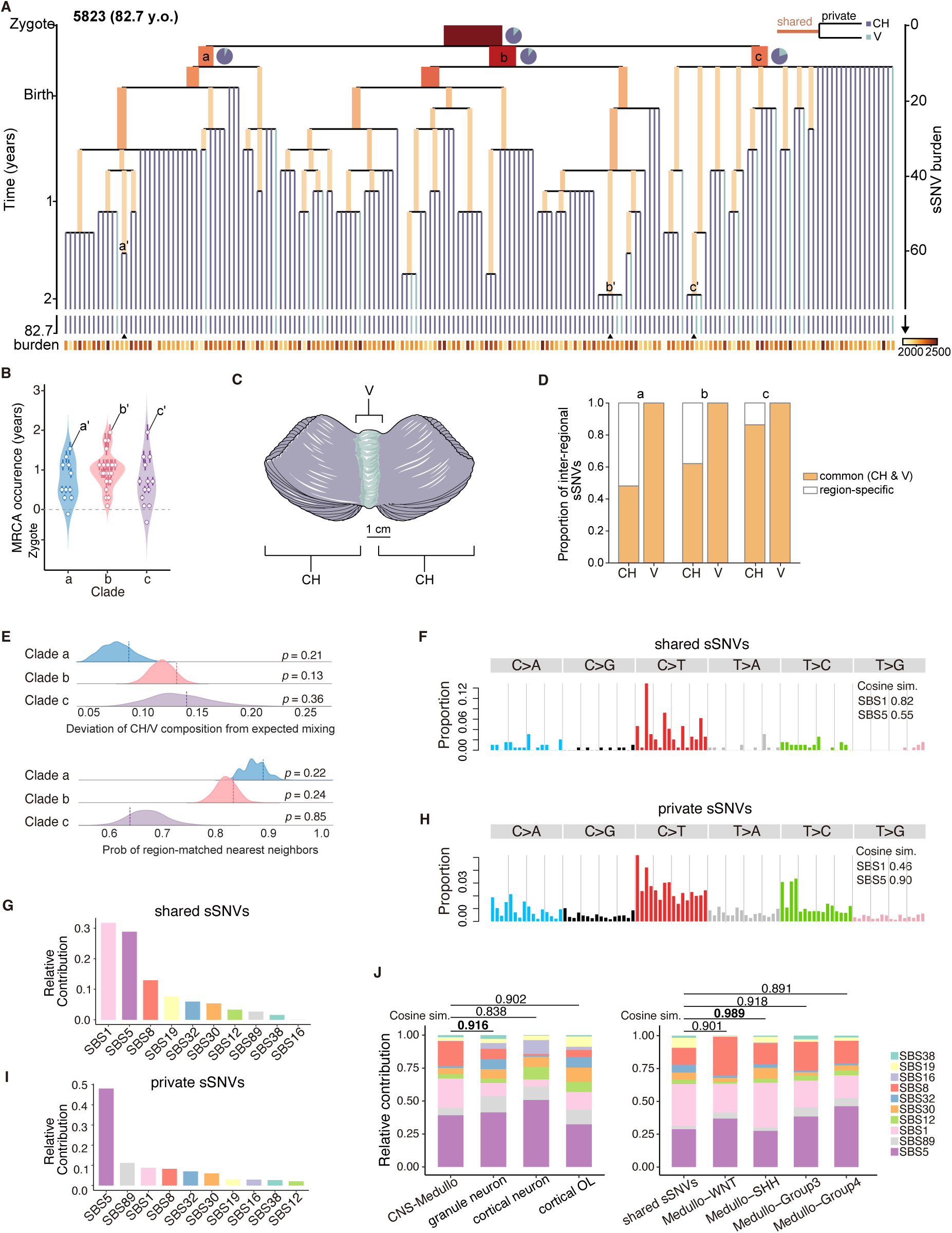
Single cell lineage tracing of cerebellar GNs. A. Early clonal phylogenies reconstructed for subject 5823 (82.7 year old; 178 cells). Tree depth reflects the number of accumulated sSNVs. Orange branches represent shared sSNVs; purple and light-blue branches depict private sSNVs unique to CH or V, respectively. Branch lengths are proportional to the number of sSNVs; branch width corresponds to the fraction of cells carrying the sSNVs. Pie charts categorize clones based on two regions (CH and V) at internal bifurcation. The annotation (bottom) shows the per-cell sSNV burden. Triangles mark the latest-born sub-clades (a’, b’, c’) derived from each of the three major clades (a–c). The developmental timing (left) was inferred from the linear mixed-effects model. B. Violin plots showing the distribution of the estimated MRCA occurrence time (in years from zygote) for clade a–c. Each point represents an inferred ancestral clone from the shared-mutation branches, with vertical error bars indicating the 95% confidence intervals derived from the linear mixed-effects model. C. Schematic depicting CH and V of adult human cerebellum. Adapted from Servier Medical Art (https://smart.servier.com) licensed under CC BY 4.0. D. Bar plots showing the proportion of common (orange) versus region-specific (white) sSNVs between CH and V cells within clade a–c. E. Measures of anatomical intermixing in the phylogenetic tree. Top: A region-mixing deviation index was computed to quantify how strongly the CH:V composition of inferred subclusters deviates from the overall CH:V ratio of each clade. Ridge density curves show the null distributions generated from 1,000 permutations of region labels, with vertical dashed lines indicating the observed values. None of the clades exhibited significant deviation from random mixing (permutation test: *p* = 0.21 for clade a; *p* = 0.13 for clade b; *p* = 0.36 for clade c). Bottom: Distribution of the probability that a cell’s nearest neighbors originate from the same anatomical region. Ridge curves show null distributions from 1,000 label permutations; vertical dashed lines indicate the observed values. No clade exhibited significant enrichment of region-matched neighbors beyond random expectation (permutation test: *p* = 0.22 for clade a; *p* = 0.24 for clade b; *p* = 0.85 for clade c). F–I. The mutational spectrum and contributions of COSMIC SBS signatures for shared (F, G) and private sSNVs (H, I). Shared sSNVs represent variants mapped to ancestral branches of the lineage tree, while private sSNVs represent terminal-branch variants unique to single cells. J. Signature contributions in medulloblastoma versus GNs, cortical neurons and OLs (left); and in shared sSNVs of GNs versus medulloblastoma subtypes (right). Cosine similarities measure pairwise contribution resemblance.

To evaluate whether any region-enriched clones existed, we compared the proportion of common versus region-specific sSNVs between CH and V GNs. Although CH neurons showed more region-specific sSNVs (Fig. 5D), this is likely attributable to their larger representation in the lineage tree, which increases the probability of detecting hemisphere-restricted events. To account for this imbalance, we performed two permutation-based tests to assess whether any clade exhibited non-random anatomical enrichment (see methods), and both analyses showed no significant anatomical clustering across the three major clades (Fig. 5E).

Similar to our elderly 82.7 year old donor, we constructed a lineage tree of GNs from a 19.8 year old, albeit with fewer (41) cells, and identified several related clones that diverged around the time of birth or a few weeks postnatally (fig. S6, D and E). While the late birth of GNs is consistent with known timelines of cerebellar and granule cell development in humans (*16*, *17*, *20–23*), our findings suggest that widespread tangential migration of GNs or their precursors occurring late after birth can result in clones that are not restricted to anatomically defined areas of the cerebellum.

### Somatic mutation patterns in GN precursors mirror those in medulloblastoma tumors

Next, we asked if these clonal sSNVs that emerged early in life exhibited mutational signatures distinct from private sSNVs that accumulated later in GNs. We observed a pronounced enrichment of C>T mutations in clonal SNVs (Fig. 5F), with about 32% of clonal sSNVs attributed to SBS1—the largest relative contribution among the detected mutational signatures (Fig. 5G). Additional contributions from SBS5, SBS8 and SBS19 accounted for approximately 29%, 13% and 8% of clonal sSNVs respectively (Fig. 5G). In contrast, private sSNVs were dominated by SBS5 (48%), consistent with its established role as an age-related, clock-like mutational signature, and showed a reduced contribution from SBS1 (8.8%) (Fig. 5, H and I). These findings suggest that GNs experience different mutagenic forces in early development compared to aging.

We compared those somatic mutations arising in GN precursor cells to mutations occurring in cancer cells of the cerebellum to enlighten the cell of origin of cerebellar cancers. Granule cell precursors (GCPs) are the cell of origin of the sonic-hedgehog activated subtype of medulloblastoma (*42–44*), so we asked if mutational signatures observed in our GNs are present in medulloblastoma subtypes and to what extent. Using cosine similarity analysis of mutational signatures, we first found that GNs showed higher similarity (0.916) to a published medulloblastoma dataset (*45*), than cortical neurons and OLs (Fig. 5J). Then, we leveraged our lineage tree analysis of clonal sSNVs and compared mutational signatures to the subtypes of medulloblastoma. Interestingly, we found that clonal mutational signatures from GNs had the highest cosine similarity of 0.989 with the sonic hedgehog subtype of medulloblastoma (Fig. 5J), consistent with a likely origin of these tumors from GCPs. These analyses highlight the benefit of studying somatic mutations in normal cells as they can reveal shared mechanisms occurring in cancer cells, inform cell of origin of tumors, and ultimately treatment strategies for cancer.

## Discussion

Our data show that the mutational forces active in GNs show overlapping, but also some remarkably different patterns, from excitatory cerebral cortical neurons, despite GNs also functioning as excitatory neurons. Surprisingly, mutational patterns in GNs more closely resemble mutational patterns in OL glia than excitatory cerebral cortical neurons. The higher accumulation of cell-division-associated mutational signatures (SBS1 and SBS19) in aging GNs compared to cortical neurons is quite surprising, given that both neuronal types are considered postmitotic. This raises the possibility of ongoing low-frequency cell division events in the neuronal lineage in the aging cerebellum although we only found direct evidence for ongoing neurogenesis for about 2 years after birth. The absence of adult-born GNs from our 178 single cell analyses despite the accumulation of SBS1 mutations could reflect not capturing enough cells, or that GN birth past the early postnatal period does not occur. The latter case would prompt a re-examination of cellular processes driving SBS1 and SBS19 mutational signatures.

The sIndel mutation spectrum of GNs was also more similar to OLs than cortical neurons. Differences in somatic mutation rates and spectra seen in GNs may be linked to decreased transcriptional levels and complexity in GN and OLs compared to cortical excitatory neurons. Moreover, GNs have smaller nuclei and cell bodies compared to cerebral cortex excitatory pyramidal neurons (*16*, *46*), which may reflect differences in chromatin packing, organization, and possibly the amount of DNA exposed to mutation or repair. Indeed, our data suggest that somatic mutations in GNs are depleted in more accessible chromatin regions, similar to OLs, but in contrast to larger excitatory neurons where the reverse relationship holds.

Consistent with this notion is a preprint report that Purkinje cells, one of the largest neuronal types, accumulate somatic indels at twice the rate of cerebral cortical neurons (*47*). These differences suggest unexpected potential relationships between neuronal type, nuclear size, and chromatin organization to somatic mutation accumulation. Given the broad differences between GNs and cerebral cortical neurons, diverse neuronal types in the human CNS may be expected to have surprisingly diverse patterns of age-related mutation, which could have relevance for understanding the cell type-specific vulnerability so often seen in age-related neurodegenerative conditions.

The observation that CH and V neurons can originate from a single common ancestor as late as 2 years postnatally was surprising, because it indicates that daughter cells born this late postnatally must travel to different anatomical regions that could be separated by millimeter or even centimeter long distances, corresponding to hundreds to thousands of cell body lengths . It remains to be determined if there are specific regions of the developing cerebellum that harbor these late-born neurons, and if such migration is driven by the same signals that guide early-born GNs such as semaphorin-6A, brain derived neurotrophic factor (BDNF) and Bergmann glia fibers (*19*, *48*).

Our finding that clonal sSNV mutational signatures closely resemble those of the sonic hedgehog activated medulloblastoma subtype supports the suggestion of GCPs as the cell of origin of this subtype of medulloblastoma. Moreover, it highlights the fact that mutational patterns in normal cells can be used to identify cell lineages of interest in cancer initiation. Identifying the mutational signature differences between normal cerebellar cells and malignant cells may reveal biological pathways that can be targeted to treat cancers of the cerebellum including medulloblastoma. This principle more broadly can be applied beyond cancers of the brain to other cancer types.

## Acknowledgments

We thank R. Matthieu and the Boston Children’s Hospital Flow Cytometry core for assistance with nuclei sorting. We thank R. S. Hill and J. E. Neil for their administrative help, and members of the Walsh and Huang labs for fruitful discussions. We thank the NIH Neurobiobank, donors of postmortem tissues, and their families for their contributions to science. **Funding**: This work was supported by a Hanna Gray Fellowship from the Howard Hughes Medical Institute to K.E., an Alzheimer’s Association Research Fellowship to A.Y.H, a Burroughs Wellcome Fund Career Award for Medical Scientists to S.K., grants from the National Institutes of Health (NIH) K08NS128272 to S.K., R01NS032457, R01AG070921 to C.A.W, R01AG088082 to A.Y.H, R56AG079857 to A.Y.H. and C.A.W. This research is supported by the NIH Common Fund, through the Office of Strategic Coordination/Office of the NIH Director under awards UG3NS132138/UH3NS132138 and UG3NS132144/UH3NS132144 to C.A.W.. This publication was made possible through the support of Grant 62587 from the John Templeton Foundation to C.A.W. The opinions expressed in this publication are those of the author(s) and do not necessarily reflect the views of the John Templeton Foundation. C.A.W. is an Investigator of the Howard Hughes Medical Institute.

## Author contributions

Conceptualization: KE, YY, AYH, CAW

Methodology: KE, YY, AYH, CAW, EG, MM, ZA, SM, CC, LS, BF

Investigation: KE, YY, AYH, CAW, EG, CNC, MM, BF, SM, ZA, CC, LS

Funding acquisition: SK, AYH, CAW

Supervision: AYH, CAW

Writing – original draft: KE, YY, EG, AYH, CAW

Writing – review & editing: KE, YY, SK, AYH, CAW

## Data and materials availability

New sequencing data generated in this study will be deposited in a public repository, with controlled use conditions set by human privacy regulations. All scripts will be made available on GitHub. Previously published PTA data for neurotypical controls were downloaded from dbGaP (phs001485.v3.p1) and NIAGADs (NG00162).

## Supplementary Materials

### Materials and Methods

#### Human tissue sources

Frozen post-mortem brains were obtained from the NIH Neurobiobank at the University of Maryland Brain and Tissue Bank, and research on these de-identified samples was performed at Boston Children’s Hospital with approval from the Institutional Review Board (S07-02-0087). All the neurotypical control subjects used in this study had no known clinical history of neurological disease or were listed as “unaffected control” in the biobank. Most of the neurotypical subjects were also used in prior studies (*7*, *11*) including the 82.7 year old subject 5823, where no clinical neurological diagnosis was known, and a neuropathological diagnosis included cerebral amyloid angiopathy and Braak stage III Alzheimer’s Disease Neuropathologic change.

#### Nuclear isolation and sorting from frozen post-mortem brain

Fresh-frozen post-mortem human brain tissue, previously stored at -80°C, was dissociated on dry ice into a 7 mL Dounce homogenizer on ice with 1 mL of chilled nuclear isolation media (250mM Sucrose, 25mM KCL, 5mM MgCl_2_, 10mM Tris-HCl pH 8, 0.1% Triton X-100, 0.2U/ul RNase inhibitor, protease inhibitor, and 1mM DTT). 50mg of tissue was homogenized on ice and carefully layered on top of 14 mL of chilled sucrose cushion buffer (1.8M sucrose, 3mM MgCl_2_, 10mM Tris-HCl pH 8, 0.2U/ul RNase inhibitor, protease inhibitor, and 1mM DTT) in a 15 mL Falcon tube. The samples were spun in the centrifuge for 1 hour at 30,000 xg and the supernatant was then removed and discarded, leaving the pellet in about 100 μL sucrose cushion buffer. The pellet was then resuspended in 400 μL of chilled blocking buffer (0.8% Bovine Serum Albumin in 1X PBS, 0.2U/ul RNase inhibitor, protease inhibitor, and 1mM DTT), and the suspension was set on ice for 10 mins. After 10 mins, the suspension was filtered through a 40 μm filter into chilled 1.5 mL Eppendorf Lo-Bind tubes, which were then centrifuged for 10 mins at 900 xg. The supernatant was then removed, leaving 75 μL of total volume remaining. Blocking buffer was added to reach a total volume of 500 μL. The following antibody and nuclei stain were then added: anti-NeuN (Sigma Aldrich catalog# MAB377X) at 1:500, DAPI (ThermoFisher Scientific catalog# 62248) at 3:500. The antibody-stained nuclei were then incubated on a shaker at 4°C for 30 mins protected from light. The tubes were then centrifuged for 5 mins at 400 xg. The supernatant was removed, leaving 50 μL of nuclei pellet solution remaining. Blocking buffer was then added to reach a total volume of 500 μL. The nuclei pellet was resuspended, and the suspension was centrifuged and washed again at least an additional time with blocking buffer. The pellet was then resuspended and filtered through 40 μm filters into fresh, chilled 1.5 mL Eppendorf Lo-Bind tubes for sorting. NeuN+ nuclei were then identified by flow cytometry using forward scatter and side scatter gates (fig. S1A) to refine the target population. snRNA-seq as described below was performed to verify purity of the target NeuN+ populations.

#### 10x Genomics snRNA-seq

Nuclei from fresh-frozen post-mortem human brain tissue were prepared and sorted as described above. To prepare single nuclei for RNA sequencing, up to 13,000 nuclei were sorted directly into 1.5 mL Lo-Bind tubes containing Master Mix supplemented with Nuclease-Free Water, as described in the 10X Genomics Chromium Single Cell 3’ Reagent Kits v3.1 User Guide (CG000315 Rev E). The GEM generation & barcoding, post GEM-RT cleanup & cDNA amplification, and 3’ gene expression library construction were performed as per the manufacturer’s instructions. Libraries were then analyzed using the Agilent 4200 TapeStation System and sent to Psomagen for sequencing on the Illumina Novaseq X platform (2 x 150bp reads)

#### snRNA-seq analysis

Read alignment and gene expression quantification were performed using CellRanger (v8.0.1) with the human reference genome GRCh38-2024-A. Ambient RNA contamination was removed using CellBender (v0.3.0) (*50*), an unsupervised probabilistic model applied to raw count matrices to infer and subtract background noise, producing denoised expression matrices for downstream analysis. Subsequent quality control was performed using Seurat (v4.3.0) (*51*). Cells with ≤300 or ≥30,000 UMIs, ≤200 or ≥7,000 detected genes, or ≥10% mitochondrial UMIs were excluded. Genes detected in fewer than three cells were removed. Doublets were detected and excluded using scDblFinder (v1.16.0) (*52*). Expression values were normalized and log-transformed using “Seurat NormalizeData”, scaled using “ScaleData”, and dimensionality reduction was performed by principal component analysis on the top 2,000 variable genes selected using the “vst” method. Batch effects were corrected with Harmony (v1.2.3) (*53*), and clustering was performed using the Louvain algorithm via functions “FindNeighbors” and “FindClusters”. The final cell types were annotated based on published cell type-specific gene markers (*27*, *28*).

#### Single-cell whole genome amplification and sequencing

NeuN+ nuclei from frozen post-mortem brain tissue were isolated as described above. Single nuclei were then sorted into individual wells of a 96-well plate containing BioSkryb PTA Cell Buffer. PTA was performed largely according to the manufacturer’s protocol (ResolveDNA Whole Genome Amplification kit P00001-07292022) but with the following modifications. Briefly, cell lysis (addition of MS Mix to cells) was performed by mixing the plate at 1400 rpm for either 20 mins at room temperature (RT) per manufacturer’s protocol, or 30 mins at 4°C to improve yield. After addition of the SN1 and SDX, the samples were incubated at RT up to 20 mins. Amplification with the reaction mix was performed as described in the manufacturer’s protocol. Post-amplification products were then cleaned using a 2X bead prep using AmpureXP beads (Beckman Coulter Life Sciences catalog #A63882) and the yield was quantified using the Qubit 1X dsDNA, high sensitivity assay kit (ThermoFisher catalog #Q33231). Using single cell whole genome amplified (by PTA) product, libraries were constructed using KAPA HyperPlus Kits (Roche catalog# KK8514). See manufacture’s protocol or (Bioskryb SkrybAmp for Ultra-low DNA Inputs; High Performance DNA Amplification from Small Samples P/N 100068). PTA product planned for 30X sequencing were sent to Psomagen Inc. for library preparation. For samples planned for 10X sequencing, libraries were prepared similarly. Briefly, no fragmentation was performed. The End Repair and A-Tailing reaction mix of Ultrapure Water, Fragmentation Buffer, ER/AT Buffer, ER/AT Enzyme was added to 300-500ng of PTA products in a final volume of 35ul using nuclease-free water. The End Repair and A-Tailing was then performed by incubating the samples at 65°C for 30 mins. To each sample, KAPA UDI Adapters (Roche catalog# 08861919702) were then diluted to 5µM using KAPA Adapter Dilution Buffer (Roche catalog# 08278539001). These adapters were then added to each sample along with nuclease-free water, Ligation Buffer, and DNA Ligase. This was incubated at 20°C for 15 mins to ligate the adapters. Post-ligation products were cleaned using a 0.8X AmpureXP bead prep and 10X KAPA Library Amplification Primer Mix and 2X KAPA HiFi HotStart Ready Mix were added directly to the bead slurry prior to amplification. Samples were amplified with 10 PCR cycles and cleaned up using two consecutive 0.55X→0.8X double sided bead preps using AmpureXP beads. Quality of completed libraries were assessed using the Agilent 4200 TapeStation System. Libraries were then sequenced by Psomagen Inc. on Illumina Novaseq6000 or NovaseqX platforms (2 x 150bp).

#### Single-cell multiplex PCR and analysis

Quality control assay on amplified genomes was performed using a multiplex polymerase chain reaction (PCR) of four random genomic loci as previously reported (*8*, *12*, *54*). Briefly, 1μL of amplified post-PTA product was added to 20 μL of a PCR master mix containing 0.05 U/µL Phusion Hot Start II DNA Polymerase, 1µM Multiplex Primer Mix, 500uM dNTPs, and 1.25X Phusion HF Buffer. The reaction mixture was then placed in a thermocycler programmed to the following conditions: 94°C for 15 mins, 13 cycles of [94°C for 1 minute, 68°C for 1 minute with -1°C/cycle, 72°C for 1 minute], 35 cycles of [92°C for 1 minute, 55°C for 1 minute, 72°C for 1 minute], 72°C for 10 minutes and a 4°C hold. The multiplex PCR products were analyzed using a QIAxcel Advanced System as per the manufacturer’s instructions and samples that showed 4 discrete bands <1000 bp were used for further analysis.

#### Bulk genomic DNA extraction and sequencing

For newly generated bulk genomic DNA sequencing data, DNA extraction was performed on frozen post mortem human brain tissue using the DNeasy Blood & Tissue Kit (Qiagen catalog# 69504) as per the manufacturer’s instructions.. Genomic DNA was sent to Psomagen Inc. for library preparation and sequencing.

#### Alignments for bulk and PTA data

BWA (v0.7.15) (*55*) was used to align the bulk WGS and PTA scWGS reads to the human reference genome (GRCh37 with decoy) to generate BAM format files. Picard Tools MarkDuplicates (v2.8.0) was run with duplicates marked. Indel realignment and recalibration of base quality scores were performed using Genome Analysis Toolkit (GATK) (v3.5) (*56*).

#### Quality control of scWGS

Sequencing depth was calculated from aligned BAM files using samtools (v1.15.1) (*57*). The evenness of single-cell genome amplification was evaluated using two metrics: median absolute pairwise difference (MAPD) and coefficient of variation (CoV). MAPD quantifies local amplification variability by computing the median absolute difference between log2-transformed copy number ratios of every pair of neighboring bins constructed with equal numbers of uniquely mapped reads. CoV captures genome-wide amplification variability and was calculated as the ratio of the standard deviation to the mean of bin-wise copy number ratios. Higher values of either metric indicate increased amplification unevenness.

#### Somatic mutation calling from PTA data

Single Cell ANalysis 2 (SCAN2, v1.1) (*8*) was used to detect sSNVs and sIndels from single-cell PTA data with matched bulk data. A cross-sample panel was first constructed using the “scan2 makepanel” with the following parameters: human reference genome GRCh37 with decoy sequences (--ref), dbSNP v138 common variants (--dbsnp), 1000 Genomes Phase 3 SHAPEIT2 phasing reference panel (--shapeit-refpanel), and metadata mapping each single-cell ID to its corresponding individual (--makepanel-metadata). Somatic mutation detection was subsequently performed on a per-individual basis using “scan2 call_mutations” with the following parameters: each PTA single-cell BAM file (--sc-bam), the matched bulk BAM file (--bulk-bam), and the cross-sample panel (--cross-sample-panel). For each cell, we obtained SCAN2 variant allele fraction (VAF)-based sSNV and sIndel calls, estimates of mutation detection sensitivity, and autosomal genome-wide burden from the corresponding scan2_object.rda file. Next, mutation-signature-based rescue was performed using “scan2 rescue” with the parameters: --rescue-target-fdr=0.01, and single-cell scan2_object.rda. Rescued mutation calls were used in detection of shared mutations and phylogenetic tree construction.

#### Linear mixed-effects modeling of mutation burden

To quantify age-associated accumulation rates of somatic mutations, we used the linear mixed-effects regression models from the lme4 (v1.1.36) R package. Both genome-wide and signature-specific mutation burden were modeled as continuous response variables. P values for pairwise comparisons of slopes were computed with the emmeans R package (v2.0.0), which utilizes Tukey-adjusted t-tests. To test the age effect of mutation burden in cortical neurons, cortical OLs and GNs from neurotypical individuals, we fitted the model *y_ijk_* = (*β_j_* + *μ_ij_*) × *ρ_i_* + *ε_ijk_*; *ρ_i_* = *α_i_* + *c*, where *y_ijk_* is the mutation burden in cell *k* from cell type *j* of individual *i*, *β_j_* is the fixed-effect of age in cell type *j*, *μ_ij_* is the individual-specific random effect on the age slope for cell type *j* following a normal distribution with mean 0 and variance *σ*, *ρ_i_* is the chronological age of individual *i* measured from fertilization, *α_i_* is the age of individual *i* at birth, *c* = 268/365 (268 days, median ovulation-to-birth interval (*58*)), *ε_ijk_* is the cell-level residual measurement error following a normal distribution with mean 0 and variance *τ*. To obtain QC-corrected mutation burden estimates, we extended the mixed-effects model by including an additional covariate, *θ_ijk_*, representing the QC metric (MAPD, CoV, sequencing depth, and mutation detection sensitivity), thereby re-estimating the age effect while accounting for technical variability.

#### Mutational signature analysis

To mitigate overfitting during mutational signature decomposition, we applied forward stepwise regression to preselect COSMIC v3.2 signatures (78 SBS and 18 ID signatures), as described previously (*59*). Signature inclusion was iteratively evaluated based on improvements in non-negative least squares fits, quantified by reductions in the sum of squared error (SSE) computed using the lsqnonneg function from the pracma R package (v2.4.4). The stepwise procedure was terminated when the SSE improvement fell below 30,000 for SBS signatures or 2,000 for ID signatures. This approach resulted in the selection of 10 SBS and 8 ID signatures, which were then refitted using MutationalPatterns (v3.14.0). Signature-specific somatic mutation burden was obtained by multiplying the contribution by the corresponding genome-wide mutation burden.

#### Annotation of genomic location and functional categories

Genomic annotations for sSNVs and sIndels were assigned using ANNOVAR (version updated in 2025March2nd) (*60*) based on the Func.refGene field. Variants were first classified into intergenic or genic categories. Genic variants were further subdivided into upstream (within 1 kb upstream of the transcription start site), 5′ UTR, exonic (coding sequence excluding untranslated regions), 3′ UTR, downstream (within 1 kb downstream of the transcription end site), splice-site (within intronic 2 bp of a splicing junction), and intronic regions. Variant counts were aggregated by cell type. To ensure robust estimation and avoid instability from sparse categories, only annotation categories with at least 20 total sSNVs or sIndels in any of the three cell types were included in the enrichment analyses shown in Fig. 4A–D. sSNV and sIndel functional consequences were annotated using SnpEff (v5.0) (*61*). Functional impact levels (High, Moderate, Low, and Modifier) were assigned using the first entry in the ANN field of SnpEff-annotated VCF files.

#### Permutation of mutation calls

Permutation-based background mutation sets were generated to control for potential biases introduced by trinucleotide context and the distribution of phaseable regions. In each permutation, the mutation calls were randomly shuffled within phaseable regions of the genome on a per-cell basis, while maintaining chromosome assignment and trinucleotide context. We performed 1,000 permutations of the sSNV and sIndel list for each cell using “scan2 permtool” with the parameters: the mutation calls (--permtool-muts), each single-cell ID (--permtool-sample) and 1000 permutations (--permtool-n-permutations).

#### Mutation enrichment analysis

We assessed enrichment and depletion of sSNVs and sIndels across genomic regions defined by gene expression, replication timing, and chromatin accessibility. Gene expression profiles for GNs were derived from in-house snRNA-seq data, whereas profiles for cortical neurons and OLs were obtained from Jeffries et al. (*49*) Replication timing data were obtained from ENCODE Repli-seq across 15 cell lines (*62*). Chromatin accessibility data for GNs were derived from a published human cerebellar snATAC-seq study (*28*), with corresponding profiles for cortical neurons and OLs sourced from our prior work (*11*).

For gene expression enrichment analysis, genes were ranked according to their expression levels and partitioned into five equal-sized expression groups. sSNV and sIndel densities were calculated separately for each group. Expected mutation densities were estimated using permutation sets (see “permutation of mutation calls”). Observed and expected counts were aggregated at the individual level, and enrichment ratio was calculated as the ratio of observed to expected mutation counts for each expression group. This analysis was performed separately for each cell type.

Replication timing signals were imported from bigWig files and partitioned into five equal-sized groups based on replication timing scores, ranging from early- to late-replicating regions. sSNV and sIndel densities were calculated for each replication timing group. For each cell type, individual-level enrichment ratios were first computed separately for each Repli-seq cell line in the same way as above and subsequently summarized by taking the median across the 15 cell lines.

For chromatin accessibility enrichment analysis, accessibility tracks of GNs generated in hg38 were converted to hg19 using UCSC LiftOver to ensure coordinate consistency across datasets. Accessibility signals were aggregated into non-overlapping 1kb genomic bins with UCSC bigWigAverageOverBed. Genomic bins were classified into five accessibility groups, with bins showing zero signal designated as group 1 and bins with detectable accessibility divided into four equal-sized groups (2–5). Mutation enrichment ratios were calculated as described above.

#### Mutational signature enrichment analysis

Signature-specific enrichment analysis was conducted using the 10 COSMIC SBS signatures identified above (see “mutational signature analysis”). sSNVs overlapping each genomic group (e.g., gene expression, replication timing or chromatin accessibility group) were refitted to the selected signatures using MutationalPatterns fit_to_signatures function. Permuted mutation sets were processed identically to obtain expected signature contributions. Signature-specific enrichment ratios were calculated as the ratio of observed to expected mutation counts for each signature within each genomic group.

#### Gene Ontology analysis

Gene Ontology enrichment analysis was performed on genes with high-impact indels using GOseq (v1.54.0) (*63*) after controlling for gene length bias. GO terms with P < 0.05 are reported.

#### Detection of clonal sSNVs

For 30X PTA data, we used rescued sSNV calls generated by SCAN2. To increase sensitivity for detecting low-frequency somatic variants, we applied more lenient germline filtering criteria in matched bulk WGS data, including ≤1 low-quality alternative read (max.balt.lowmq), ≤ 10 total alternative reads (max.bulk.alt), and VAF ≤ 0.3 (max.bulk.af).

For 10X PTA data, a subset of variants was found to recur across multiple cells within the same batch, indicative of batch-related technical artifacts. To mitigate these artifacts, VAF-based sSNV calls were first aggregated across all cells within 10X PTA batch. For every candidate variant, mutation status was re-inferred from the raw BAM file of each cell using the Bayesian genotyper MosaicHunter (*64*) to estimate posterior probabilities of mosaic genotypes. Variants were subsequently projected into a matrix between recurrence ratio and 96-trinucleotide context, where recurrence ratio was defined as the proportion of cells within a batch harboring the variant. Variants exceeding a recurrence ratio threshold of 60%, informed by the recurrence ratio-context distribution, were classified as technical artifacts and excluded from rescued sSNV calls with lenient germline filtering criteria as described above.

For 250X bulk WGS data, sSNVs were called using MosaicForecast (*41*). Variants with mosaic_P ≥ 0.6 were retained as high-confidence calls, and variants overlapping gnomAD (*65*) were removed to exclude potential germline variants.

Finally, sSNVs identified from 30X PTA, 10X PTA, and bulk WGS data were integrated on a per-individual basis. Clonal sSNVs were defined as variants detected in at least two single cells, or in at least one single cell and a bulk sample. These clonal variants were re-evaluated using MosaicHunter in the same way as above to confirm the mutation status. To further reduce false positives introduced by lenient germline filtering, variants were filtered out if they showed excessive recurrence (recurrence ratio > 0.6) or insufficient allelic support (median VAF ≤ 0.4 across mutant cells). High-confidence clonal sSNVs were then assembled into a variant–sample (cell and bulk) genotype matrix per individual shown in fig. S6B–C.

#### Phylogenetic tree construction

Phylogenetic trees were constructed for each individual through four steps. (1) Generation of the variant-cell genotype matrix. The variant-cell genotype matrix was extracted from the results described in “Detection of clonal sSNVs”. The shared sSNVs among cells were kept for phylogenetic analysis. (2) Inference of clonal structure. To identify groups of sSNVs that co-occur across cells (hereafter referred to as mutation modules), the genotype matrix was subjected to hierarchical clustering using the pheatmap R package with complete linkage and binary distance. This step was used to reveal structured co-occurrence and mutual exclusivity patterns among sSNVs, providing an initial partition of sSNVs into nested candidate clone-defining sets. Variants incompatible with tree-like inheritance were subsequently pruned.

Specifically, sSNVs were removed if they displayed random distributions across cells without forming stable co-occurrence patterns, or generated conflicts with nested inheritance, such that their presence-absence patterns could not be embedded into a hierarchical clonal structure. (3) Construction of phylogenetic tree. A lineage tree was then assembled by organizing the retained mutation modules into a hierarchical structure based on set inclusion relationships across cells. The tree was rooted at the earliest ancestral node, corresponding to the clone defined by the shared sSNVs present in the largest fraction of cells. Parent-child relationships between nodes were determined by the nested structure of mutation modules. Branch attributes were defined from the genotype matrix and the inferred hierarchy: branch lengths indicate the number of sSNVs acquired along that lineage; branch widths reflect the fraction of descendant cells carrying the corresponding sSNVs, providing clone prevalence within the cell population. (4) Developmental timing estimation and tree annotation. The shared sSNV count was extrapolated to a genome-wide burden after correcting for mutation detection sensitivity. These burdens were then used to infer the timing of ancestral events using the linear mixed-effects model (see “linear mixed-effects modeling of mutation burden”). Anatomical region labels were overlaid onto the phylogenetic tree for visualization.

The time to MRCA was calculated as the average inferred developmental time from zygote across cells. To assess whether cells from the CH and V exhibited non-random mixing in the lineage tree, we quantified the deviation of regional composition from that expected under random mixing. For each clade, we first calculated the global proportion of CH cells, and for each ancestral clone within the clade, we computed the absolute difference between the clone-specific CH proportion and the clade-level CH proportion. These differences were weighted by clone size and summed to generate a region-mixing deviation index for each clade. Statistical significance was assessed using 1,000 permutations in which regional labels were randomly shuffled among cells, and the observed deviation index was compared with the permutation-derived null distribution. We also estimated the probability that a cell’s nearest neighbors arose from the same anatomical region. Nearest neighbors were defined as cells within the same ancestral clone. Region-matching probabilities were computed for each cell and averaged to obtain a clade-level metric. Statistical significance was assessed in the same way as above.

## Supplementary Figures

**Fig. S1.**
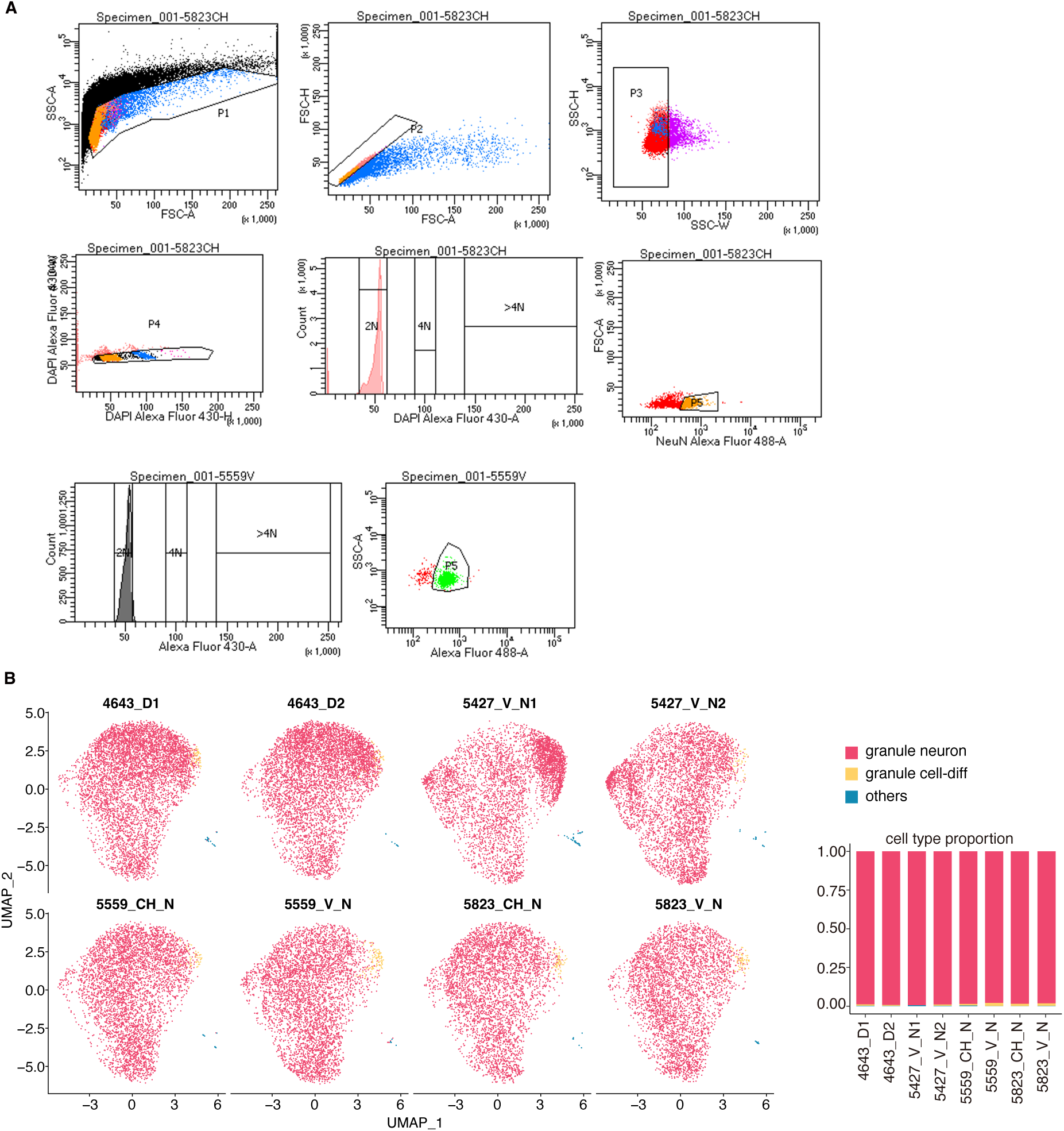
FANS of neuronal nuclei from cerebellar brains and validation of purity. A. FANS showing forward scatter and side scatter gates, DAPI+, 2N nuclei, and NeuN+ (P5) gates. B. UMAP plots of snRNA-seq profiles (left) and cell-type composition (right) across samples.

**Fig. S2.**
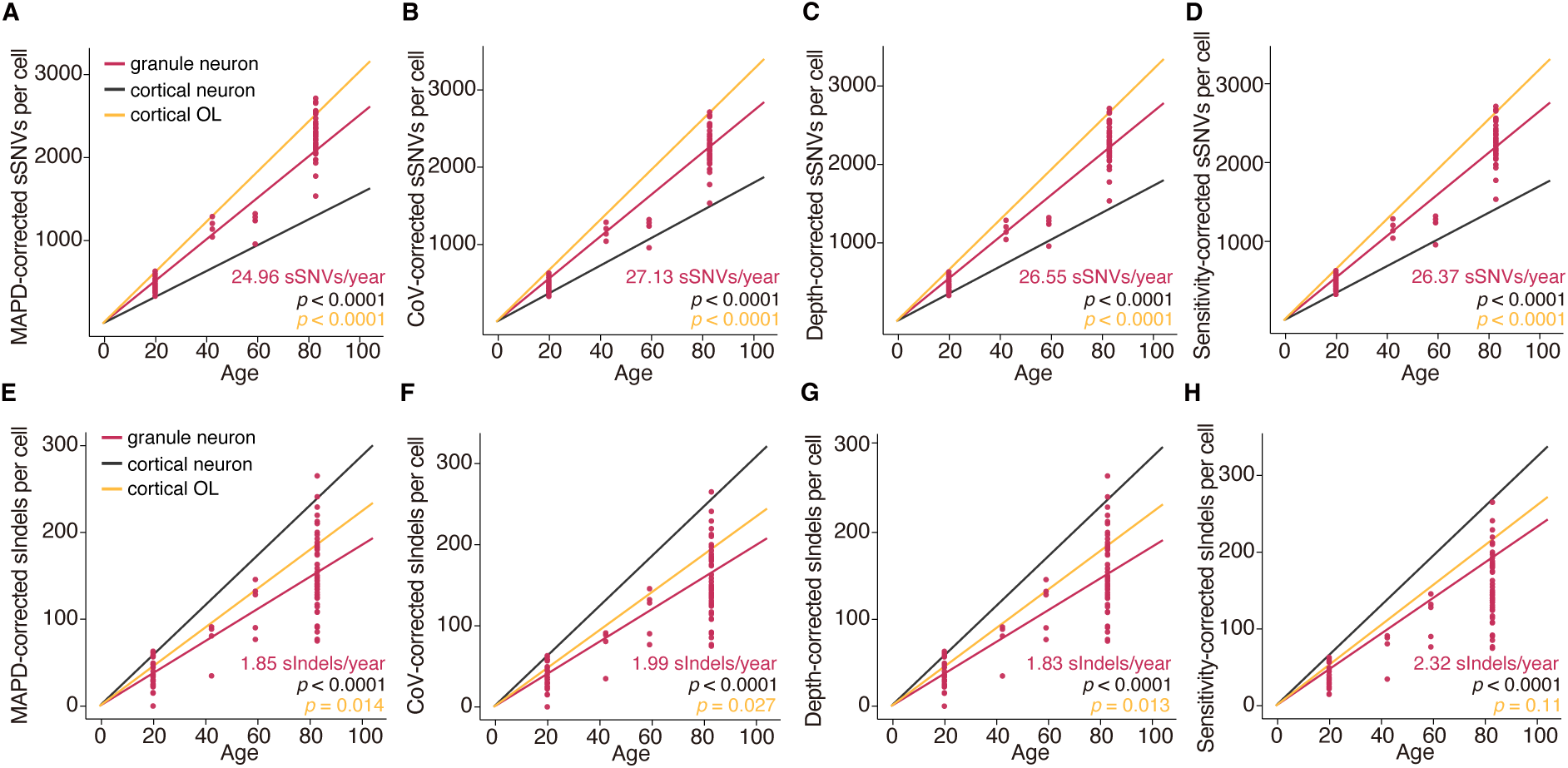
sSNV and sIndel burden comparison after controlling for QC metrics. A–H. sSNV (A–D) and sIndel (E–H) burdens in GNs, cortical neurons and OLs after correction for MAPD (A, E), CoV (B, F), sequencing depth (C, G), and SCAN2 mutation detection sensitivity (D, H). P values compare GNs with cortical neurons and OLs.

**Fig. S3.**
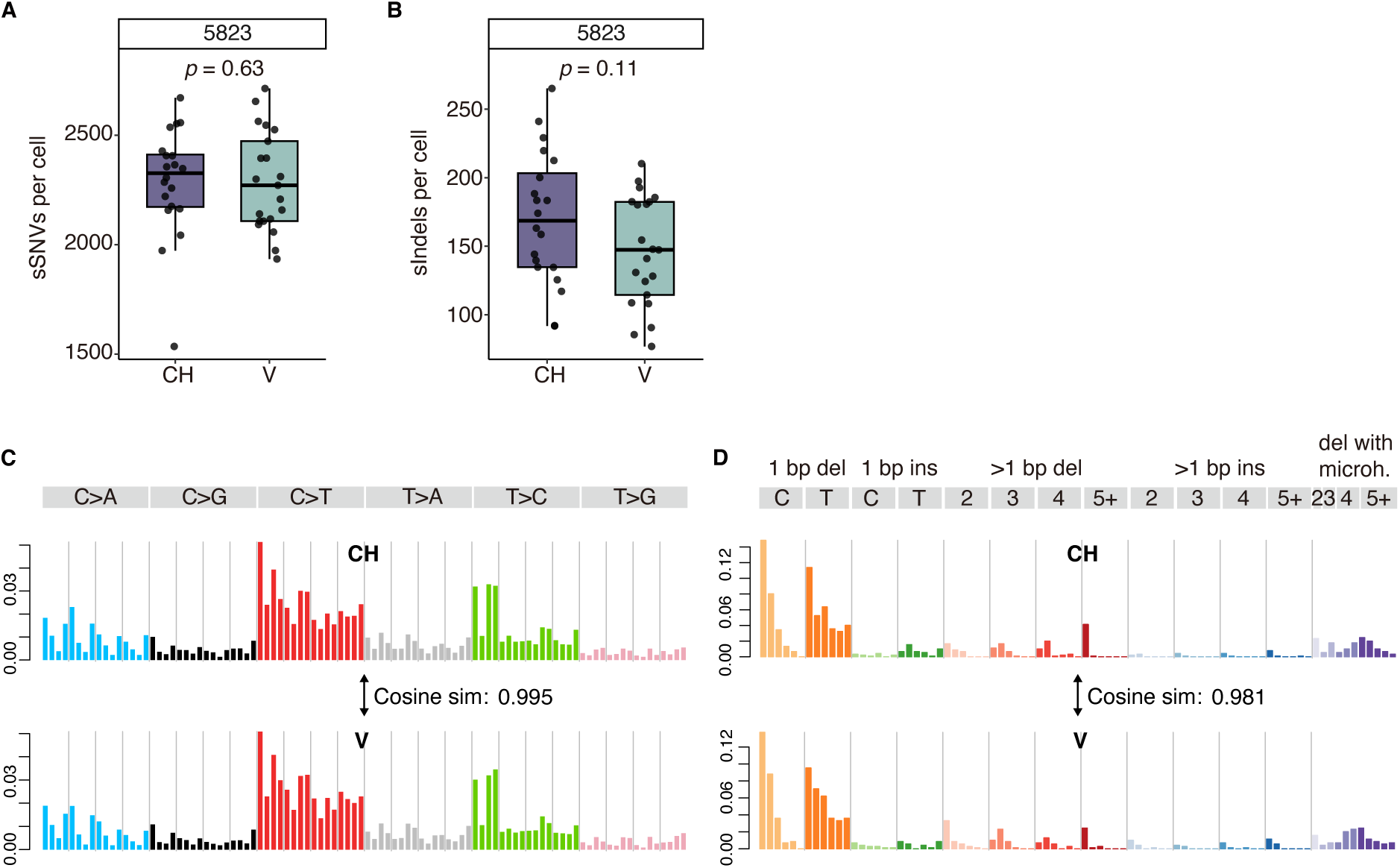
GNs from CH and V show comparable somatic mutation profiles. A–B. Comparison of sSNV (A) and sIndel (B) burden in GNs from the cerebellar hemisphere (CH) and vermis (V). P values are computed using two-tailed Wilcoxon tests. C–D. Substitution (C) and indel (D) spectra of CH and V GNs. The text in the figure indicates the cosine similarities between regional profiles.

**Fig. S4.**
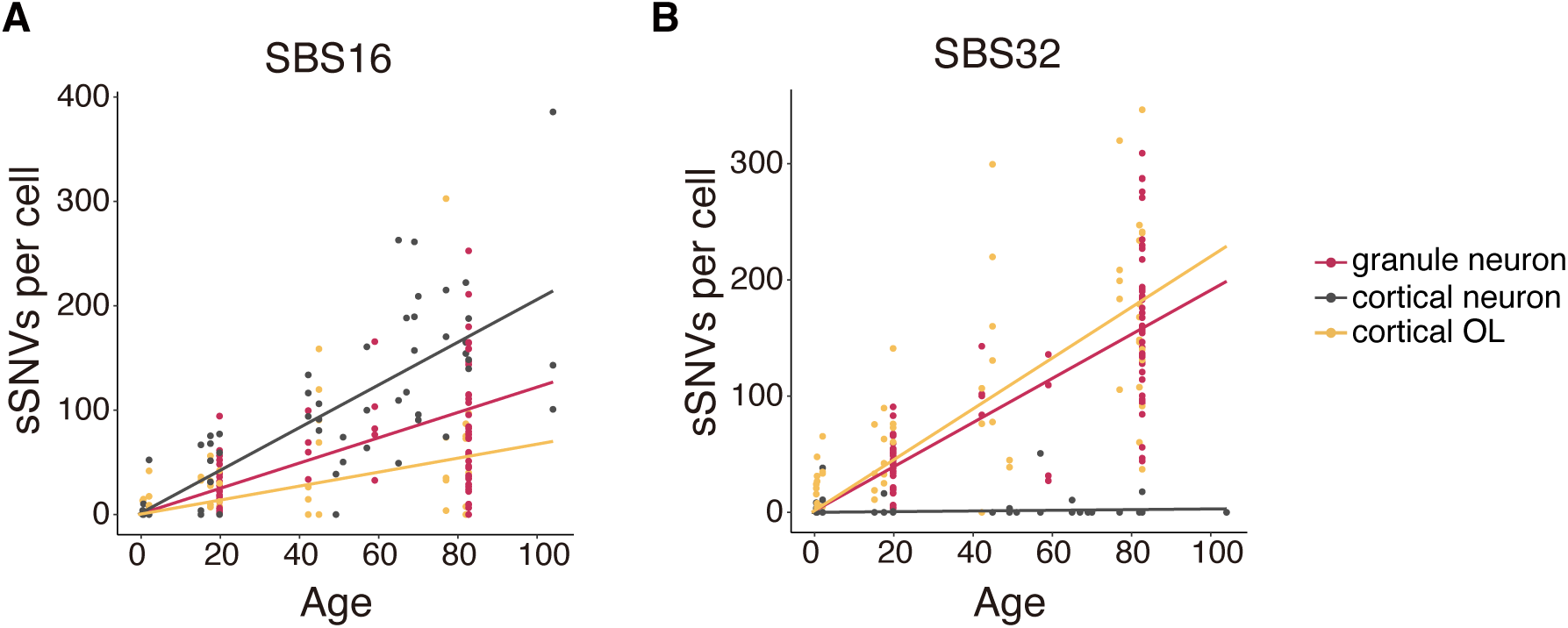
Signature contributions across three cell types. A–B. Signature contributions of SBS32 (A) and SBS16 (B) in GNs, cortical neurons, and OLs plotted against age.

**Fig. S5.**
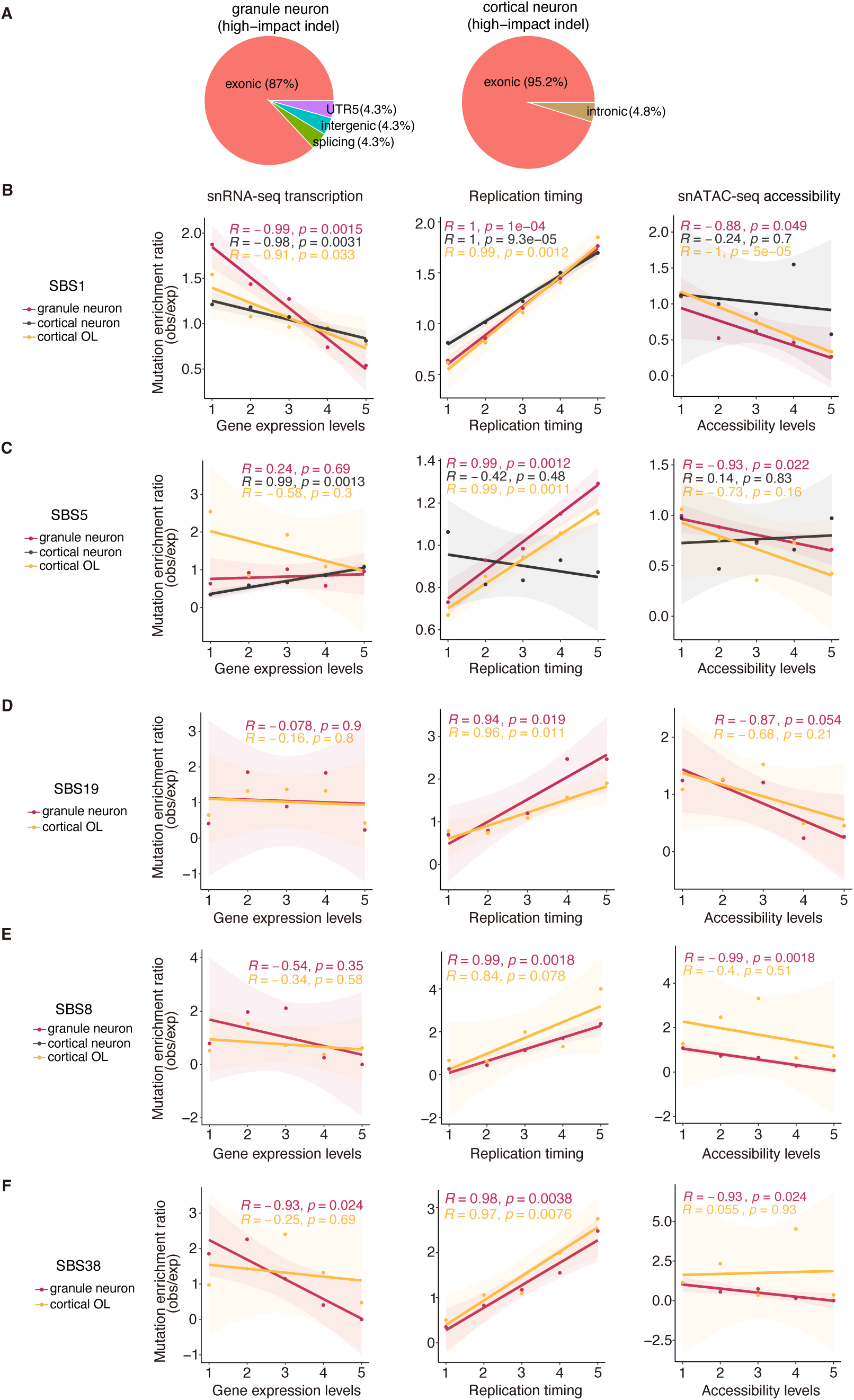
Enrichment analysis of SBS signatures in GNs. A. Pie charts showing the genomic distribution of high-impact indels identified in GNs (left) and cortical neurons (right). B–F. sSNV enrichment for SBS1 (B), SBS5 (C), SBS19 (D), SBS8 (E) and SBS38 (F) in relation to gene expression, replication timing, and chromatin accessibility.

**Fig. S6.**
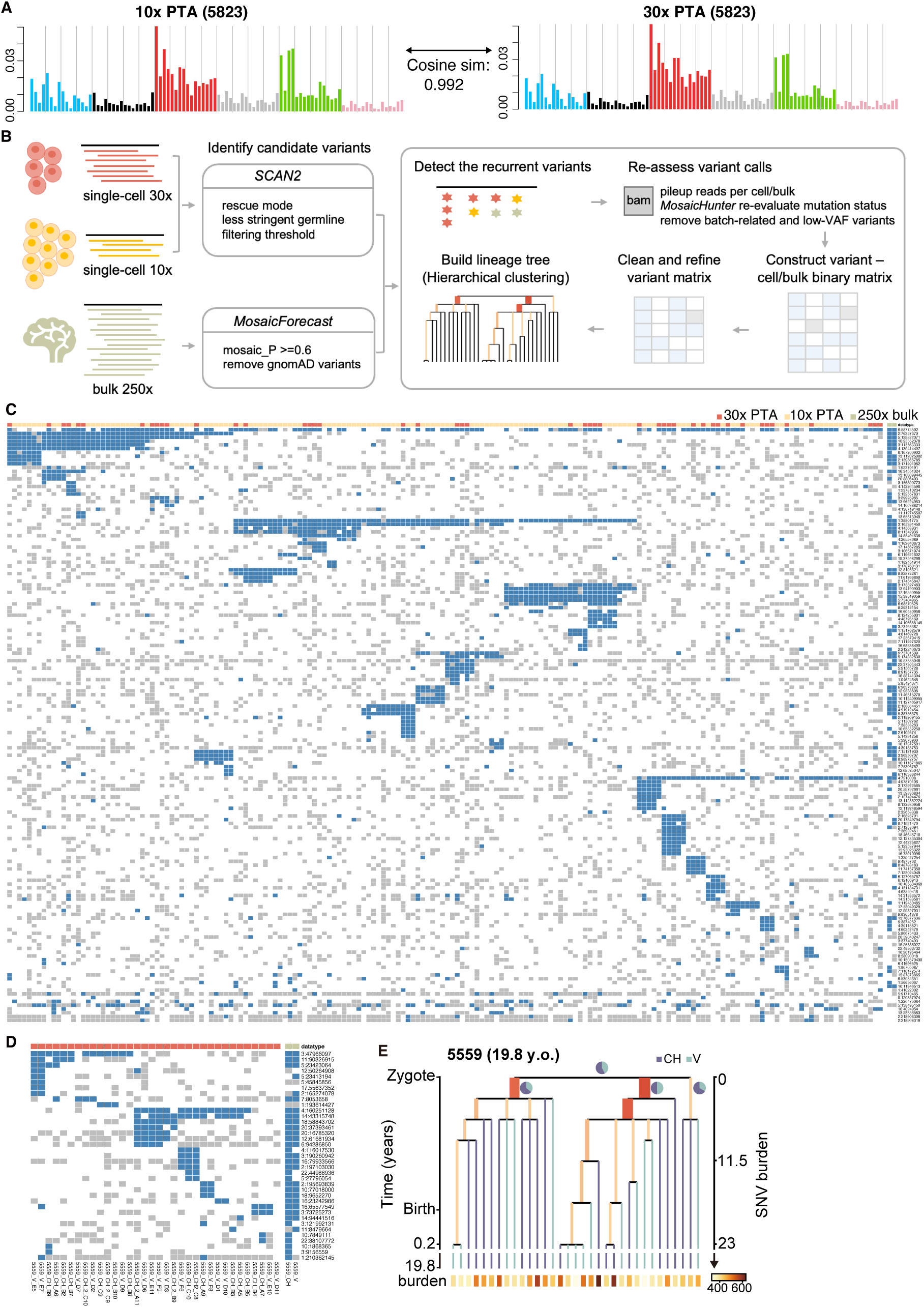
Lineage tree analysis of GNs. A. sSNV spectra of GNs from 30X and 10X PTA. B. Schematic overview of the variant-calling and lineage reconstruction workflow. sSNVs from single-cell genomes (30X PTA and 10X PTA) were identified using SCAN2 with rescue mode and lenient filtering, while sSNVs from 250X bulk WGS were called using MosaicForecast. For each subject, variants from single-cell and bulk data were merged to detect recurrent mutations, which were subsequently re-evaluated per sample (single cell or bulk) using MosaicHunter to refine mutation status and remove batch-related and low-VAF artifacts. A variant-sample binary matrix was then constructed (blue: mutant; white: wild-type; grey: undetectable due to insufficient coverage) and cleaned to eliminate potential artifacts inconsistent with lineage structure. Hierarchical clustering was applied to reorder the matrix and generate the corresponding lineage tree. C-D. Heatmaps showing the variant-sample binary matrices for subject 5823 (B) and 5559 (C), Each column represents a single cell or bulk (30X PTA: red; 10X PTA: orange; bulk 250X WGS: light green), and each row corresponds to a variant. Blue: mutant; white: wild-type; grey: undetectable due to insufficient coverage. Clonal blocks emerge as contiguous clusters of shared variants across cells, reflecting the underlying lineage structure. E. Early clonal phylogenies reconstructed for subject 5559 (19.8 year old; 34 cells). CH, cerebellar hemisphere. V, vermis (vermis cells for this experiment on 5559 were performed immediately after CH cells in the same experiment, and hence may represent a mixed population of CH and V cells).

